# Label-free imaging of large samples: 3D rendering and morphological analysis within histological workflows using serial block face imaging

**DOI:** 10.1101/2022.05.03.488912

**Authors:** Marine Malloci, Perrine de Villemagne, Paul Dorval, Magalie Feyeux, Stéphanie Blandin, Guillaume Schmid, Philippe Hulin, Perrine Paul Gilloteaux

## Abstract

Serial block face imaging (SFBI) is a method used to generate 3-dimensional (3D) reconstruction of a sample via serial image acquisition. Several SBFI approaches have been proposed for large samples, differing in the ability to generate contrast as well as in the nature of the detected signal. We propose a new system that detects the endogenous autofluorescence signal of paraffin-embedded samples. The sample preparation is simplified compared to other approaches, and adapted to be integrated into a routine histological preparation. More specifically, it was designed to limit reagent toxicity and to be compatible with downstream histological processing. We show the usefulness of the technique with a wide range of tissues based on the intrinsic autofluorescence signal. Optimization of quality section recovery offers the possibility to develop correlative approaches and multimodal analysis between the 3D dataset with the 2-dimensional (2D) sections. In addition, contrast and resolution of block-face images allow us to successfully perform post processing analysis and morphology quantifications. Overall, our methodology offers a simple, cost effective and rapid approach to obtain quantitative data on a large sample with no specific staining.

## Introduction

In biology, 3-dimensional (3D) analysis of large samples can improve our understanding of physiological and pathophysiological mechanisms. The relationship between function and 3D anatomical organization offers a holistic approach to understanding underlying biological organization.

Although conventional histology techniques provide high resolution images, 3D reconstruction of 2D sections from wax embedded samples is challenging and laborious^1–3^. The main difficulties of this approach are alignment and registration of the successive sections and section loss leading to incomplete information and artifacts in the reconstructed 3D volume^4,5^.

As an alternative, serial block face imaging (SBFI) systems have been developed. This technique is based on the following process. First, surface images of an embedded tissue block are captured digitally. Then, a section of fixed thickness is cut from the surface. The process is reiterated until the sample is entirely sectioned, allowing generation of digital volume data. All serial images being theoretically pre-aligned, they can be stacked together and converted into a 3D volume to explore it according to any virtual cutting plane.

Different home-made SBFI systems have proposed effective 3D reconstruction of wax embedded samples^6–9^. With regard to these systems, it seems that the critical point to obtain a quality 3D volume is to improve the contrast between the embedded sample and the surrounding paraffin. To make the SBFI systems compatible with paraffin-embedded specimens, several contrast-generating strategies have been proposed. One of them is to color only the exposed tissue at the surface of the block after each cut in order to contrast only the block surface to be imaged ^8,9^. However, this requires manually staining the block surface after each section which prevents the automation of the technique. Another solution is based on the use of opacified wax^6,7^. Indeed, due to the transparency of classical white paraffin, the block surface shows not only the cutting face of the sample but also deeper structures, which bleedthrough the paraffin block and impair the quality of the 3D reconstruction. This strategy further offers the possibility of higher throughput since it can more easily be automated. Opacified paraffin or stained sample approaches, and subsequent imaging using a digital color camera have provided interesting methods^6,8^. Another method based on autofluorescence detection^7^, an interesting native property in biology, can provide contrast to recover spatial and structural information.

Biological tissues have an intrinsic fluorescence, originating from endogenous fluorophores such as proteins containing aromatic amino acids, NAD(P)H, flavins and lipopigments. Plants contain many other fluorophores, such as chlorophylls, flavonoids and cell wall components. In addition, as the amount and the distribution of endogenous fluorophores can reflect physiological and/or pathological processes, auto-fluorescence monitoring can be utilized in order to obtain information about morphological and physiological state of cells and tissues^10^. Since autofluorescence levels depend on tissue composition, they can be very heterogeneous within the same organ/organism. These can provide information at the macroscopic scale but also on the internal structures of a biological sample. Because autofluorescence is ubiquitous in most living organisms, it can be used to study a wide variety of samples. Finally, its detection is simple to set up: it requires no sample treatment nor labeling and only requires an excitation source and a detector.

Among all the existing SBFI systems, one of particular interest is the episcopic fluorescence image capture system (EFIC) system^7^. EFIC sample preparation uses an opaque red colored wax that includes vybar and stearic acid, two additives for candles, and Sudan IV a lipophilic red stain^7^. This increases wax rigidity and opacity which facilitates sectioning and limits the native autofluorescence of paraffin in the visual spectrum. This system was mainly used for 3D reconstruction of mouse embryos based on autofluorescence collection.

Inspired by the EFIC approach, we have developed a simple and ready-to-use SBFI technique that does not require specific labeling or contrast agent and does not interfere with histological protocols used routinely. Sample preparation, which has been simplified compared to the EFIC method, is limited to low levels of toxicity colored paraffin embedding and is compatible with a large variety of samples. Importantly, we present a configuration adaptation to achieve better section quality and easier slice retrieval for downstream histological processing. Indeed, the system has been designed to be particularly adaptable and easily integrated onto different microtomes, as well as on cryostats. The process is controlled by a proprietary software that synchronizes block sectioning and image capturing. Volume rendering of the generated image stack provides sufficient signal-to-noise ratio to enable volume segmentation and data quantification.

## Results

### System description

The system was developed to offer an easy to install imaging apparatus, compatible with a wide variety of microtomes. It is composed of a surrounding aluminum frame, a fibered 470 nm LED as excitation source, optics to collect fluorescence signal above 500 nm and a camera. A proprietary software automates the acquisition and synchronizes the microtome rotation and the image acquisition. Custom-developed scripts can be used to run experiment-specific acquisitions.

More specifically, the sample is mounted on the microtome, and the camera is placed orthogonally to the surface of the block. The x,y, and z positions of the camera are adjusted using micrometic screws, and the focus is achieved and controlled in real time on the software interface (Figure 1). When the microtome is triggered, the motorized clamp that holds the block moves vertically towards the blade and cuts the block by coming down onto the blade. The x,y position of the block remains relatively constant throughout the acquisition as the idle position of the microtome after each successive section is fixed. This is of interest in our approach as it limits the need for post processing registration of the images. Stability of the whole system is further controlled by placing the microtome and the imaging apparatus on an optical table. This provides a mechanical reference to limit potential vibration that may occur while cutting. Optical pathways are similar to a conventional wide-field fluorescence microscopy setup. Excitation and detection have been chosen to image the autofluorescence signal. Indeed, autofluorescence is usually considered as a common drawback in light microscopy. In our approach, we rely on intrinsic autofluorescence of the tissue as the main source of signal rather than considering it as an obstacle. This has the advantage of offering a label-free approach with thus limited sample preparation. With this in mind, we chose to excite the sample with 470 nm light where the main peak of autofluorescence occurs, and detection is done above 500 nm (Figure 1d). Excitation power and integration time are controlled via the interface.

**Figure 1:**
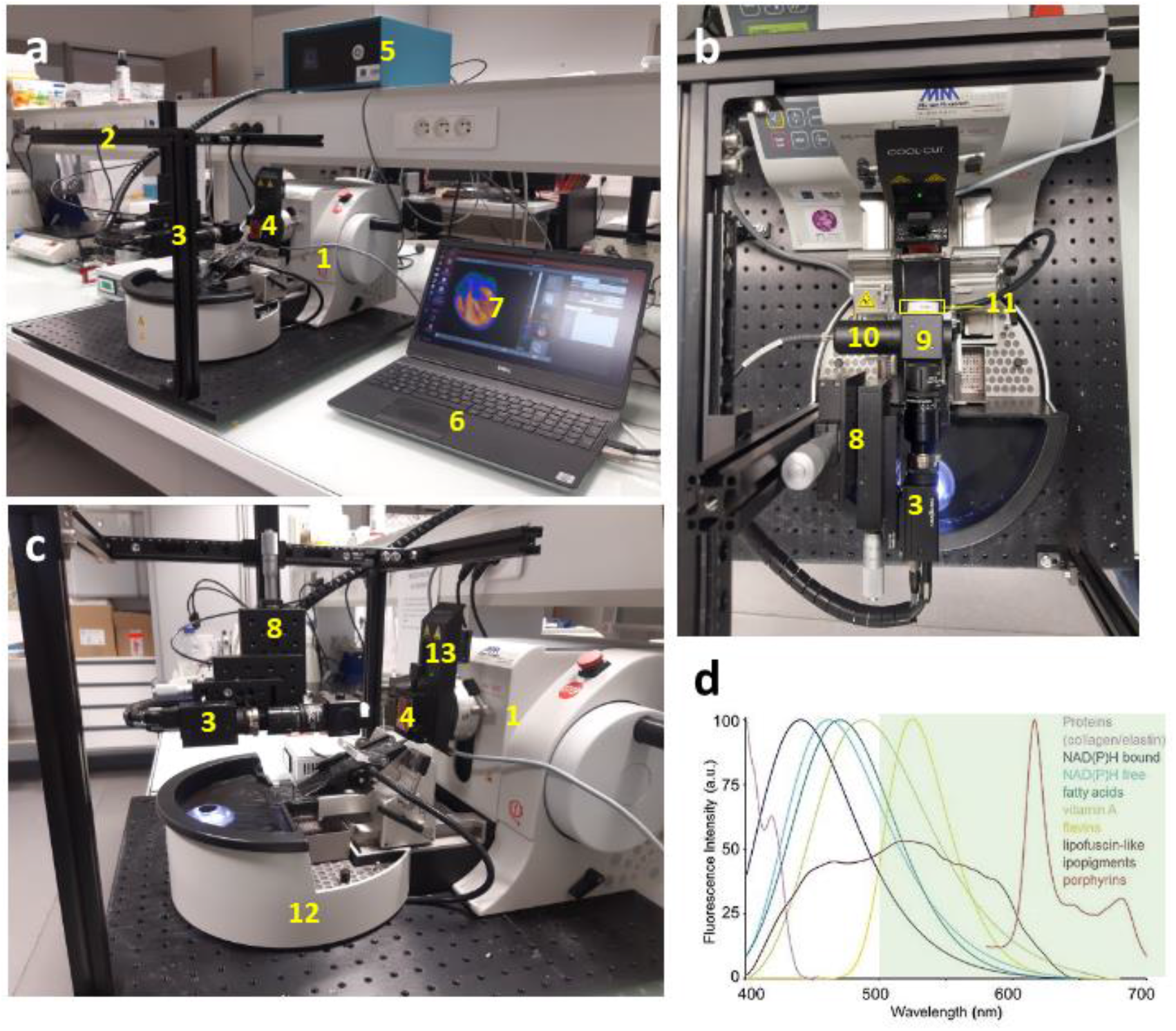
The SBFI system uses a rotary microtome with a section transfer system (STS). **(a)** Whole view image of the imaging system. The rotary microtome (1) is placed on an optical table to which is attached an aluminum frame (2) that holds the camera (3) orthogonally to the surface of the mounted sample (4). The microtome is connected to the control unit (5) which itself is connected to a laptop computer (6). The acquisitions are controlled by the control software (7) that triggers the microtome through the control unit (5). Images are stored on the laptop computer. **(b)** Top view image of the optical setup of the system. The camera is attached to Y and Z positioning stages (8). A dichroic cube is mounted on the front of the camera (9) from which the LED excitation is laterally attached (10) and onto which individual lens can be mounted to vary the field of view (11). **(c)** Lateral view showing the STS of the microtome that uses a warm laminar flow to bring cut sections into a water bath for section collection (12). The cool-cut system (13) improves section quality by cooling the block during sectioning. **(d)** Spectral profile of autofluorescence emission of individual endogenous fluorophores. Collection is done above 500nm and is shown. Graph adapted from Croce & Bottiroli^32^.

Section thickness is set on the microtome. For 3D rendering acquisitions, the thickness is set as to ensure a near-to isotropic voxel and typically varies between 3 and 20μm. When histological processing is needed, the automated acquisition is paused and sections are collected in the region of interest. Typical acquisition time for a single section is about 10 seconds. With a slice thickness of 10μm, 360μm of tissue can be processed within an hour. Thus, by considering the sample preparation steps (fixation, dehydration and paraffin embedding) and its acquisition, a 3D rendering can be obtained with this system in a maximum of 3 days.

We explored a wide range of samples to illustrate the compatibility of the sample preparation and the ease of implementation regardless of the nature of the specimen. Mammalian organs (mouse) were imaged (Figure 2a-f) as well as non-mammalian organisms such as a shrimp (g) and a zebrafish (h). The ubiquitous nature of autofluorescence allows us to also explore plants as illustrated by the broccoli (i). From these images, morphological information can be collected from both external structures (overall shape and size) as well as internal structures (organs, vasculature).

**Figure 2:**
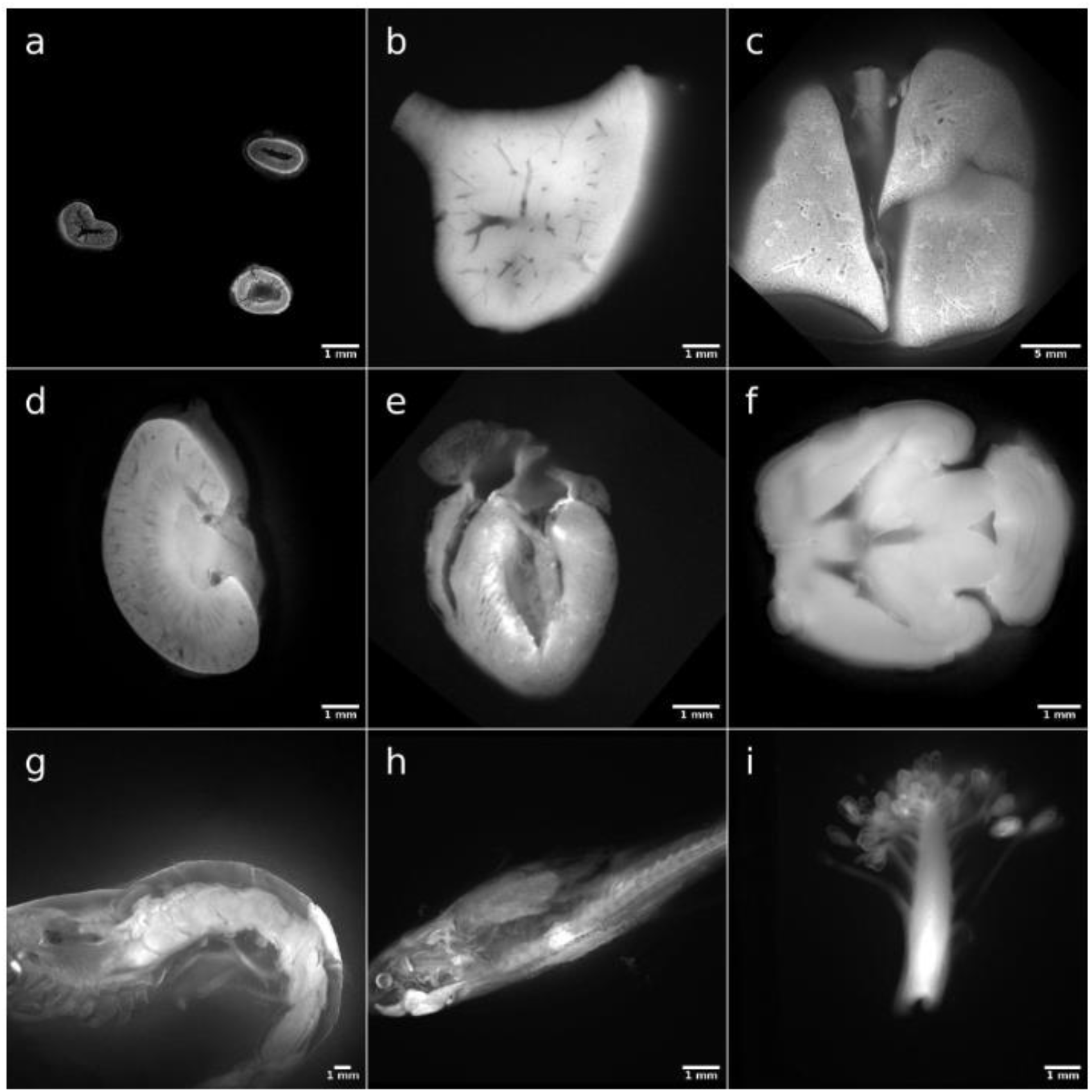
Autofluorescence signal image gallery. Acquisitions were performed on a large panel of mouse organs **(a-e)**: **(a)** intestines, **(b)** liver, **(c)** lungs, **(d)** heart, and **(e)** brain. Images **(g-i)** show non-mammalian and plant samples: **(g)** shrimp, **(h)** zebrafish and **(i)** broccoli.

We also show the acquisition of a frozen heart from mouse UBI-GFP embedded in OCT with our system associated with a cryostat (CM1950, Leica Biosystems, Nussloch, Germany). We were able to obtain autofluorescence images with sufficient contrast to distinguish internal structures (Supplemental Figure S1).

### Optimization of paraffin opacification

Typical paraffin used in histology is clear in color. As a result, the block is relatively transparent, and block surface imaging will result in unwanted recording of a large depth of field. This will negatively impact the quality of the 3D reconstruction. To enhance paraffin opacity and limit the out-of-focus fluorescence signal, we have developed a red colored paraffin to embed samples. Classic clear paraffin (Paramat Gurr, VWR) was colored by Oil red O (ORO) solution (Diapath, Italy). This lipophilic red stain is miscible in paraffin and does not interact with dehydrated tissue. Ready-to-use ORO solution was mixed at 5% volume/volume ratio (% v/v) with classic clear paraffin to produce a dark red mixture (Figure 3a).

**Figure 3:**
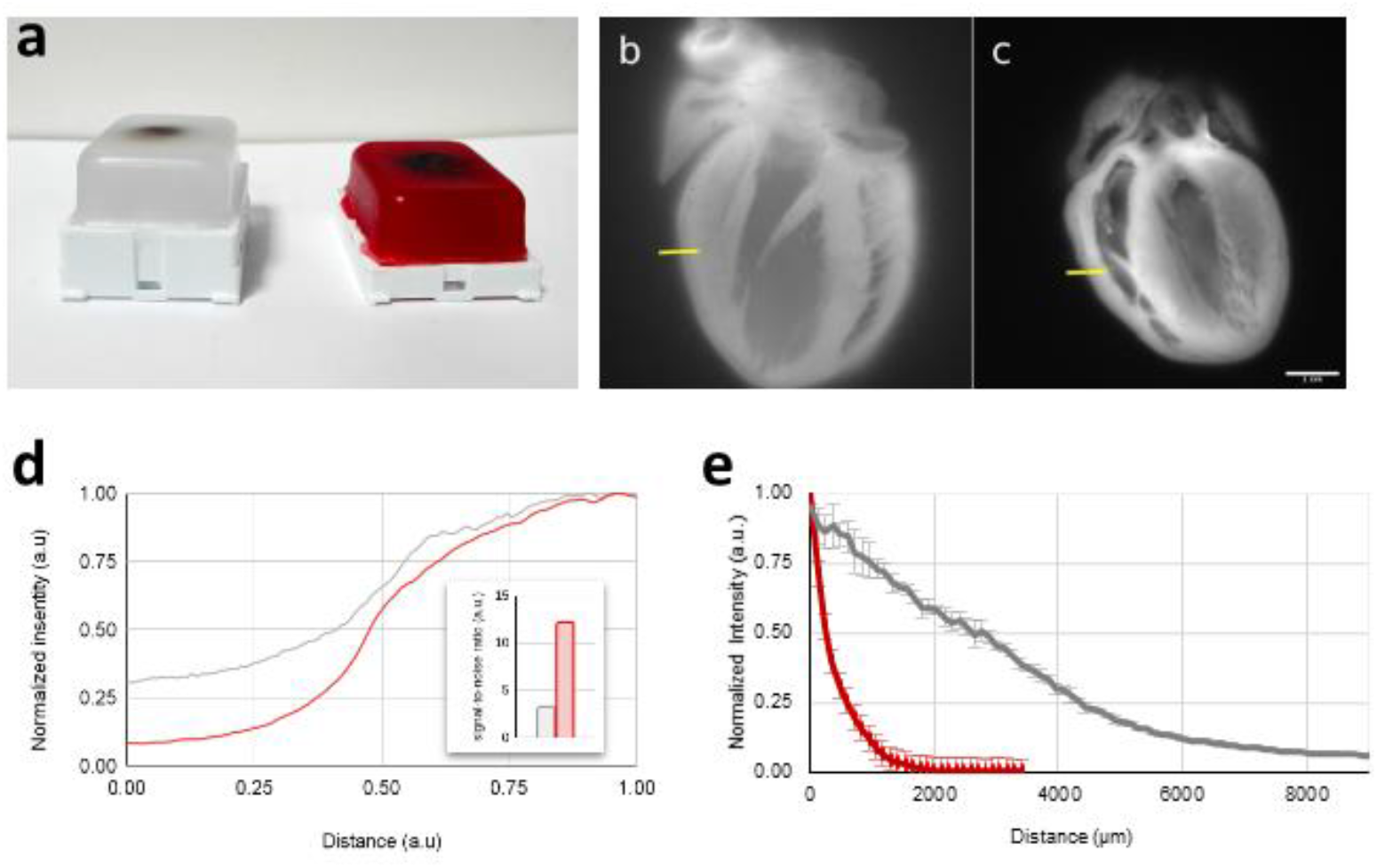
Optimization of paraffin opacification. Samples were prepared in conventional clear paraffin or in 5% ORO paraffin **(a)**. Comparison between intensity signal of rat heart embedded in clear paraffin **(b)** and in 5% ORO paraffin **(c)** were carried out. Intensity profile was normalized to the maximal value **(d)** measured on images **(b)** and **(c)** on a segment of interest (yellow line). The ratio between maximal and minimal intensity is shown in the histogram (insert). Excitation penetration in ORO paraffin was measured with comparison to clear paraffin at 470nm **(e)**. A fluorescent phantom was embedded at 45° from the surface of the block in clear (gray curve) or ORO paraffin (red curve) and intensity decay was measured on an image of the block surface. Normalized intensity profile is shown (n=6).

Rat hearts were embedded in clear and ORO paraffin and images of the corresponding block surface are shown in Figure 3b and c respectively. To show the effect of paraffin opacification on contrast improvement, the intensity profile of a segment at the sample-paraffin interface (Figure 3b and c, yellow segment) was plotted. Intensity was normalized to the maximal value. Results show that paraffin coloration reduces signal diffusion from out-of-focus tissue as seen by a net decrease in the baseline intensity of colored paraffin (Figure 3d). As a consequence, images have an improved contrast at the interface between the sample and the paraffin (Figure 3d, insert). This result was further illustrated by a measure of excitation penetration in the paraffin block in clear vs ORO paraffin. A strand of fluorescent thread was embedded with a 45° angle from the surface of the block. Then, the decay in intensity was measured accordingly as a x displacement in the recorded image which corresponds directly to the z sample depth (Figure 3e). Results show that background signal detection is reduced by 10 fold in opacified ORO paraffin compared to clear paraffin. Overall, these results show the advantage of paraffin opacification using our method to significantly improve contrast and image quality.

### Sample dimension and resolution

The samples are prepared using histology cassettes to mount the paraffin block. The maximum dimension of the samples to be studied is thus dictated by the size of these cassettes which are about 3 x 4 cm in the x,y plane. The thickness of the block is limited by the height of the mold used during the embedding, as well as by the displacement of the microtome (2,8mm). Within these maximal sample dimensions, and to explore a wider range of sample sizes, we adapted the setup to allow the user to change the field of view (FOV) of the acquisition and thus the pixel size. Individual lenses can be mounted in front of the camera to focus on smaller regions of interest or smaller samples. Using a USAF1951 resolution test chart (R1DS1P, Thorlabs), we quantified the size of the FOV for zoom factors of 0, 1, 2 and 3 and measured values of 24mm, 11mm, 7mm and 4mm respectively. Pixel size in x,y consequently varied and reached values of 23,4μm, 10,7μm, 6,7μm and 3,9μm. Results are illustrated in Figure 4 and specifications are summarized in Table 1.

**Figure 4:**
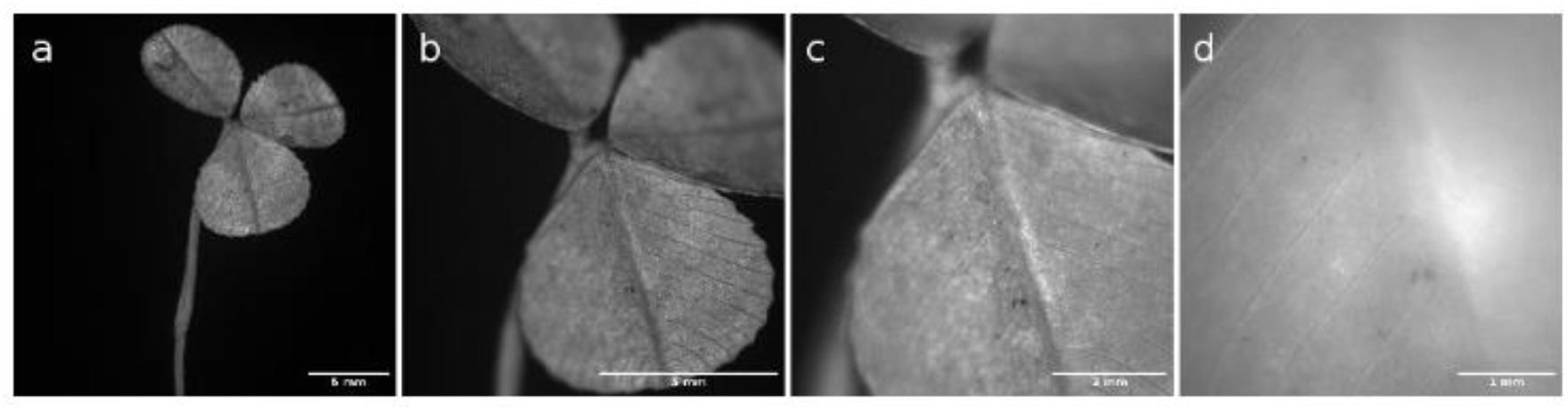
Variable fields of view. Variations of the pixel size and field of view at different magnifications allowed by the system are shown using a clover at (a) zoom 0, (b) zoom 1, (c) zoom 2, (d) zoom 3.

**Table 1:**
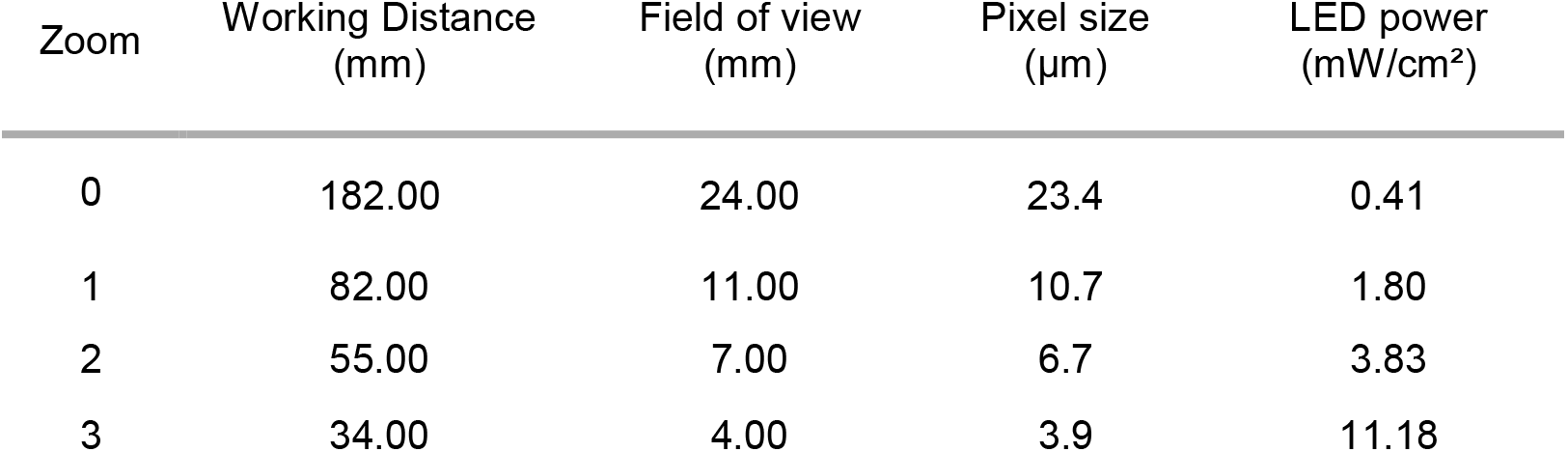
Summary of different zooms

Given the opacification properties of ORO paraffin presented previously, spatial resolution in the z axis will be dependent on the nature of the biological sample: a tissue with many cavities filled with ORO paraffin will have better resolution in the z axis than a dense sample in which the ORO paraffin will be less efficient to opacify the out-of-focus planes of the sample.

### Improved recovery of histological sections

One of the advantages of associating the imaging apparatus with a microtome is to recover sections and thus offer the possibility to do multimodal imaging. Indeed, individual sections can be collected in specific regions of interest during the acquisition. Sections can then be used for histological or immunolabeling processing. We initially adapted the system on a rotary microtome (RM2255, Leica Biosystems, Nussloch, Germany) (Supplemental Figure S2). With this setup, we were able to recover sample slices, but the section quality was not reproducible and were sometimes unusable. Indeed, it was almost impossible to recover quality sections for crumbling tissues such as the liver or bones. This is illustrated in Figure 5c for a mouse tibia section stained by Hematoxylin Phloxine Saffron (HPS) that shows poor sample integrity and debris. In addition, for these particular samples it was not possible to recover 3μm thick sections, a thickness conventionally used in histology. Rather, we were limited to 10μm sections. Successful sectioning at 3μm of this type of samples on a microtome usually requires block cooling to avoid possible deformation of the paraffin. Moreover, to help section unfolding, the user usually blows on the section to accompany the cut. This approach is no longer possible when slicing is automated during a SBFI acquisition. The possibility to recover the sections for any given sample was an important requirement in our development.

**Figure 5:**
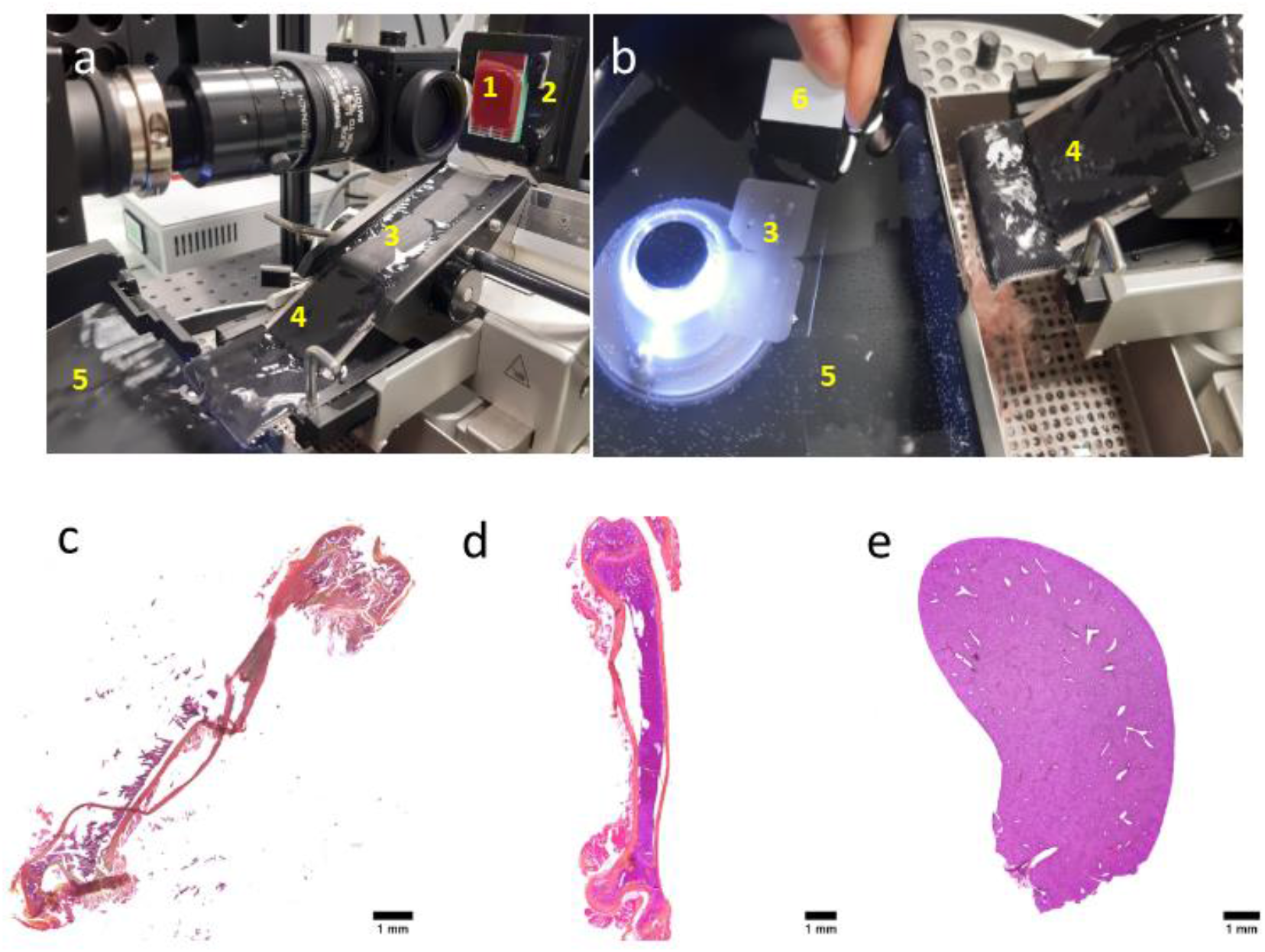
Optimization of sections recovery. The imaging setup is mounted on a microtome with a STS and cooling block. Overview of the laminar flow (a). The paraffin embedded sample (1) is sequentially cut by the microtome. Automation of the process leads to the generation of a section ribbon (3) which is unfolded by the hot water flow (4). Continuous cooling of the paraffin block by the cool-cut system (2) also improves section quality. (b) Sections (3) are manually recovered on glass slides (6) in the warm water bath (5). HPS staining of a mouse tibia section (10μm thickness) obtained on classical automatic microtome (RM2255, Leica) (c) and on a microtome with a STS and block cooling (d) (10μm thickness). HPS staining of recovered section from a mouse liver could be achieved with this approach (3μm thickness) (e). Corresponding images obtained with the SBFI system are shown in Supplemental Figure 3.

To overcome these limitations, we mounted the imaging system on a microtome with a section transfer system (Microtome HM 355S, Epredia, NH, USA) (Figure 4a). We added a cooling block (Cool-cut, Epredia, NH, USA) on the microtome rack with Peltier effect to generate a homogeneous and continuous cooling of the block throughout the cut. The section transfer system (STS) uses a hot laminar water flow to unfold sections without user help (Figure 5a). The sections are carried along by the water flow into a warm (37°C) collection bath where they can easily be collected on a glass slide (Figure 5b). This workflow greatly improves the reproducibility and quality of the sections as shown with the mouse tibia example (Figure 5c-d). Furthermore, high quality sections of 3μm thick of mouse liver could now also be recovered (Figure 5e) which was not possible on the conventional microtome during a SBFI acquisition.

### 3D rendering and image analysis

Serial block face imaging generates stacks of images in the z-axis of the sequentially cut sample. The mechanical stability of our system generates very little variation in the position of the block after each successive cut. As a result, image registration is no longer required and 3D reconstruction of the sample volume is trivial: the output of the system is a 3D stack that can be then directly rendered by volume rendering as proposed by different software. Image segmentation can be performed on these stacks to extract information of a given structure or organ. For example, from the autofluorescence stack of a mouse liver, it was possible to segment the vascular network.

Thus, the negative contrast between the non-autofluorescent vascular lumen and the autofluorescent surrounding tissue (Figure 6a) was used to extract the vascular network using Fiji^11^ (Figure 6b). Once the blood vessels have been segmented, it was then possible to reconstruct the entire network with the Skeletonize plugin (Figure 6C). Furthermore, the total vascular volume, the branch number or even the junction number were quantified as possible parameters to evaluate the physiological state of the sample, as shown in Table 2.

**Figure 6:**
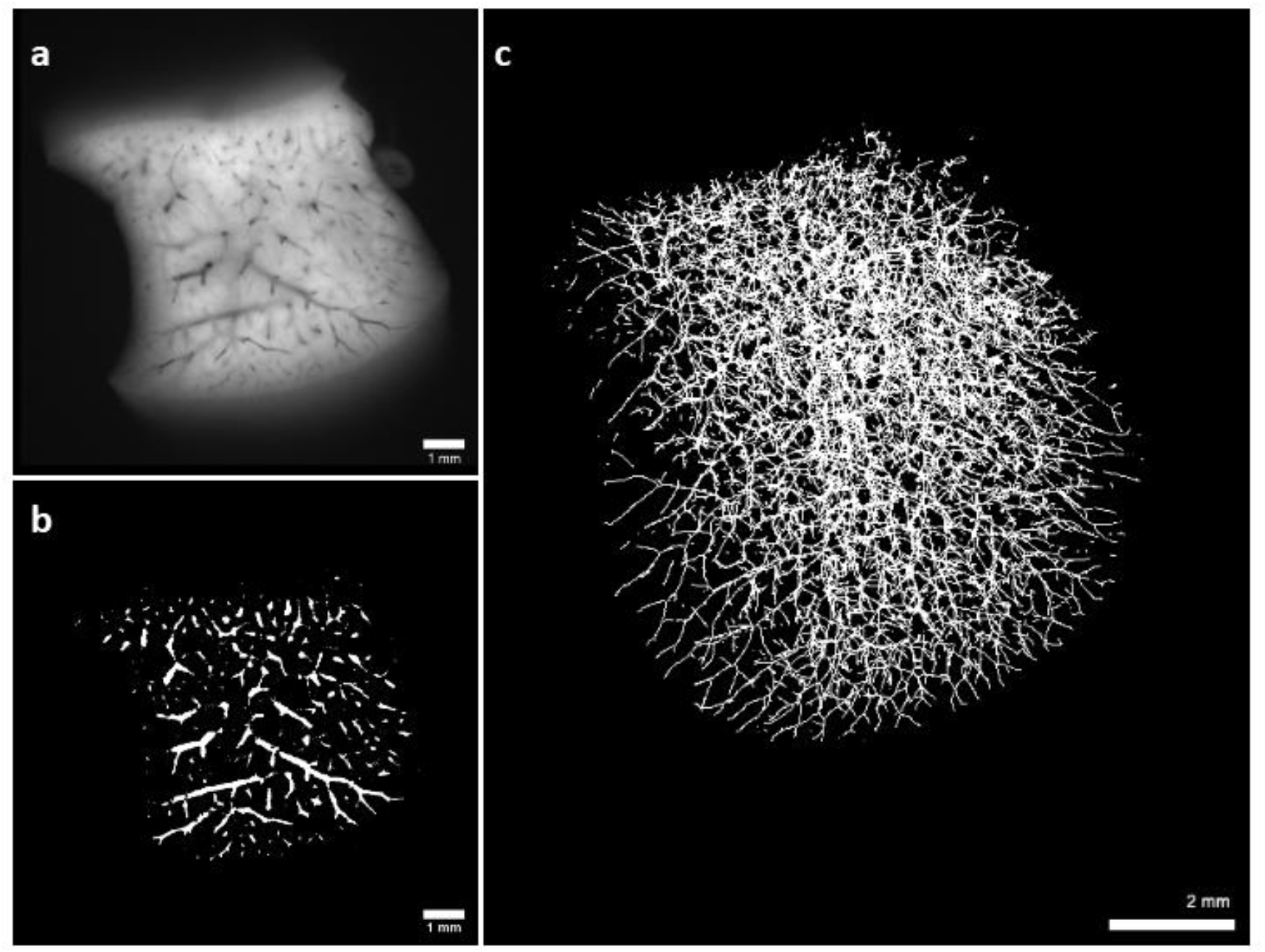
Analysis of the vascular network of a mouse liver. Autofluorescence image of a mouse liver obtained with the SBFI system: blood vessels appear black and contrast with tissue autofluorescence (a). Pixel classification was carried out using Fiji to extract liver blood vessels (b). From whole stack segmentation, vascular network was reconstructed with the Skeletonize plugin which models blood vessels in thin filaments. The network is shown in 3D rendering (c).

**Table 2:** Characterization of the vascular network of a mouse liver

| Liver Volume (cm<sup>3</sup>) | Vessel Network Volume (cm<sup>3</sup>) | Minimum Vessel Diameter (μm) | Maximum Vessel Diameter (μm) | Branch Number | Junction Number | Maximum Branch Length (μm) |
| --- | --- | --- | --- | --- | --- | --- |
| 1,089 | 0,07 | 35 | 375 | 13 618 | 6 459 | 1 432 |

### Correlative imaging

Multimodal imaging between autofluorescence images from the block face acquisition and histological stains on sections allows correlating 2D information within the 3D context of the sample. To illustrate the feasibility of multimodal imaging with our technique, we collected a mouse embryo section while the serial acquisition of autofluorescence was taking place. As such, the section can be associated with its corresponding autofluorescence image. The section was stained in Hematoxylin Phloxine Saffron (HPS) and developing organs were identified using the eMouse Atlas Project (http://www.emouseatlas.org) (Figure 7a). Using Icy software^12^, the identified organs were then bound on the stained section and transferred on the autofluorescence image obtained with our system (Figure 7b). Based on this registration between the two modalities, the individual location of the section can be visualized on the volume rendered block face acquisition (Figure 7c). Furthermore, segmentation of an individual organ such as the heart was performed on the SBFI images based on the identification performed on the HPS stained section (Figure 7d). Quantitative measurements can be performed and we typically evaluate the heart volume to a value of 7.4604.10^8^ μm^3^. With this example, we illustrate the relevance of correlative imaging using our label-free SBFI approach and histology staining to reveal internal structures and 3D organization. Both techniques, taken individually, provide a different type of information which becomes much more comprehensive when analyzed in a correlative manner.

**Figure 7:**
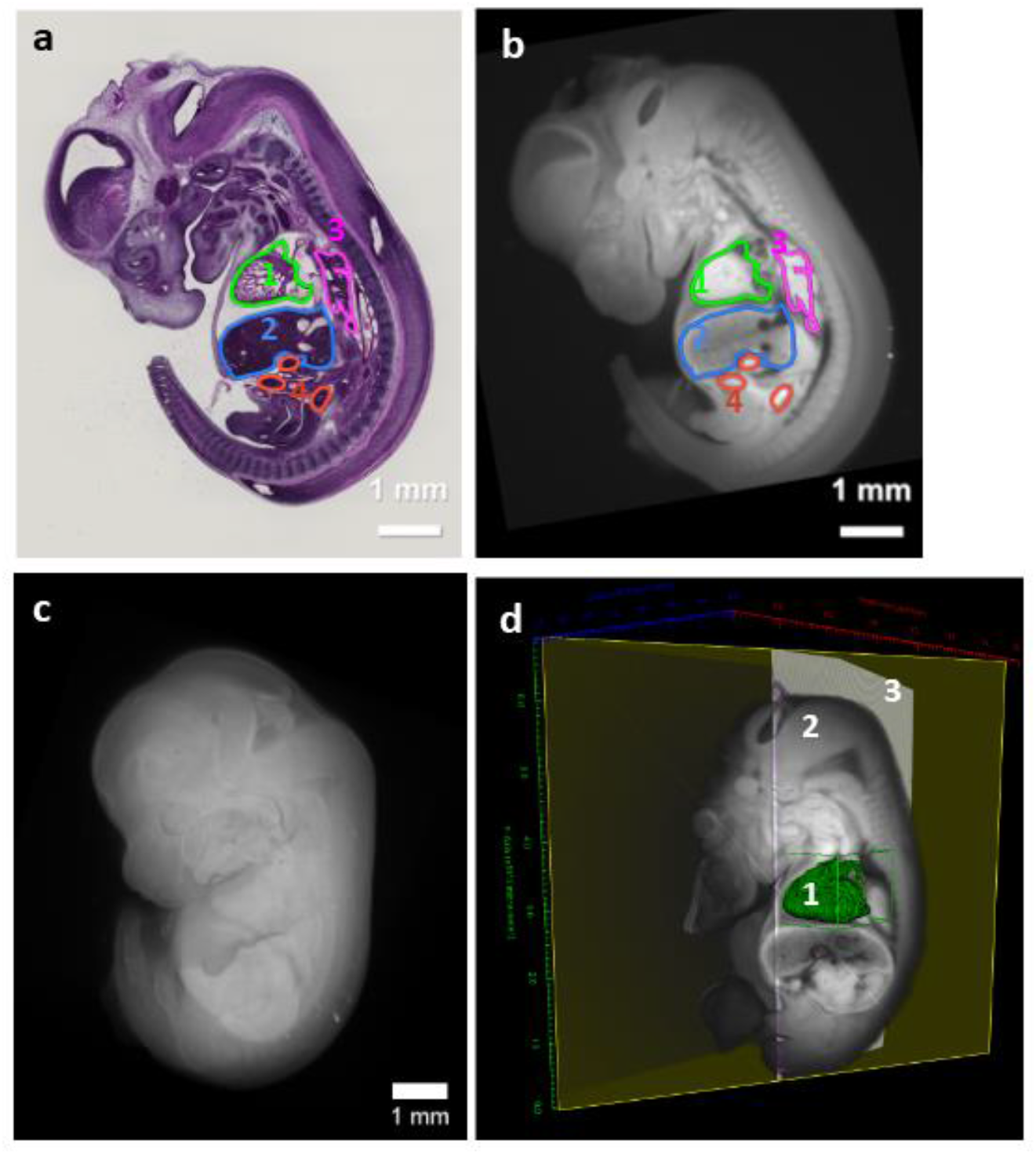
Image analysis and correlative imaging. A mouse embryo was sectioned for multimodal imaging and an individual section was collected and stained for HPS. Developing organs were identified with EMAP, manually segmented with Icy software **(a)** and transferred on the autofluorescence image **(b)** (1: heart, 2: liver, 3: lung, 4: gut). 3D rendering of embryo autofluorescence with Arivis software **(c).** 3D segmentation of the heart (1) in the 3D rendering (2) visualized using an oblique 3D slice viewer in Icy. The 2D stained section (3) was registered in the 3D volume **(d).**

## Discussion

In this report, we describe a new serial block face imaging system that detects the endogenous autofluorescence signal of paraffin-embedded samples, allowing for a rapid 3D reconstruction method. Sample preparation was optimized to enhance signal contrast. Our system offers a wide range of applications as it relies on autofluorescence which is ubiquitously expressed in the kingdom of life. It has been developed to offer an easy to install setup that uses standard cost-effective fluorescence optics, and is compatible with a wide number of commercial microtomes. We have implemented the possibility to vary the field of view and the pixel size in order to adapt the acquisition parameters to the size of the sample of interest. The acquisition is automated but nevertheless allows manual section recovery to perform multimodal imaging. Post processing image analysis and 3D volume rendering can be done. In addition, quantitative parameters can be extracted to characterize the sample.

We propose an optimized oil red-O colored paraffin to limit fluorescence bleedthrough and improve both signal to noise ratio and axial resolution. We have not observed any significant adverse effects of this staining on our ability to section paraffin blocks compared to clear paraffin, nor does it seem to interfere with the sample. We have shown that the excitation penetrates deeper into the block than the thickness of the sections performed. This, however, does not seem to affect our ability to achieve resolved 3D reconstruction. Paraffin opacification and improvement of signal to noise ratio in serial block face imaging has been addressed by others. Indeed, Ishii et al^6^ used popular white crayons to opacify the paraffin, and imaged the surface of the block with white light and a color camera. This issue was also addressed by Weninger and Mohum for the EFIC method mentioned previously^7^. In addition, they developed a similar contrast generating approach for high resolution episcopic microscopy (HREM) where resin was fluorescently doped to image the negative signal of the embedded sample^13^. All of these approaches are complementary, but have certain limitations. Indeed, the Combi approach is designed only to capture color images of samples and HREM approaches are associated with high resin toxicity and are incompatible with section recovering. The EFIC methodology, which is the most similar to ours approach, relies on the addition of a stearin and vybar mixture to opacify the paraffin, requiring higher melting temperatures. This may have detrimental effects on the sample if sections are collected to perform protein or RNA expression profiles^14^. We chose to further develop EFIC to limit modifications of the sample preparation with regards to routine protocols used in histology. In addition, we improved the setup to allow routine section collection in a region of interest during the autofluorescence acquisition for a large variety of samples. This was guided by our interest in offering a simple multimodal imaging system where 2D information can be studied within its corresponding 3D context. Ongoing work is focused on automating this process of targeting a specific region of interest for section collection by registering the ongoing acquisition in a 3D atlas or any other macro acquisition.

Imaging large and thick samples is a challenge in microscopy for optical reasons, but is relevant to study a tissue or organism as a whole. To overcome this challenge, researchers usually rely on the analysis of individual regions or subregions that are considered representative of the whole tissue or organ. Extrapolation of the information of these subsets is usually done to characterize the sample in its entirety. In comparison to existing microscopy approaches for large samples such as light sheet microscopy, the resolving power of our system remains greatly inferior (up to a few hundred nm for some developed lightsheet microscopes^15^). However, in light sheet microscopy, sample clearing is required and can be a complicated step for which the ideal conditions are sample-dependent. The reagents used are efficient for large samples but are toxic to the user, as is the case with the DISCO^16^ method which is widely used. In addition, the clearing step can require several weeks which limits the throughput of the technique. Taken together, the absence of toxic clearing reagents, the time saving protocols for sample preparation and the ease of implementation of the presented method weigh significantly among the different compromises to choose from when determining the most suited imaging approach for large samples. The resolution of our system allows us to show clear identification of both internal and external structures within a tissue (Figure 2 and 5), and illustrate a segmentation approach of fine liver vascularization (Figure 5c) and of full organ (Figure 6d). This type of information can be used to compare morphological parameters between a control and pathological state, to characterize the effect of a given drug or to evaluate developmental processes. As a proof of concept and to illustrate the quality and amount of information that can be collected solely on the autofluorescence signal, the segmented data was exported as a volume and used to generate a .stl file. Using a 3D printer, we were able to 3D print the segmented information (Supplemental Figure S4). This opens possibilities in the field of 3D bioprinting where the segmented image can be used as a template to bioprint a substrate based on true anatomical architecture using hydrogels or bioinks^17^.

Although autofluorescence of tissues provides structural information on the sample, it seems necessary to add the possibility of recovering fluorescence of specific labeling. As future perspectives, we also plan to adapt the optical path to include other excitation wavelengths to extract for example specific GFP/RFP fluorescence signals from autofluorescence. For this, paraffin sample preparation will need to be adapted because fluorophores, unlike tissue autofluorescence, are degraded during conventional paraffin embedding processes described in our material and methods. Indeed, tissue fixation and dehydration steps necessary for paraffin embedding, denature *ex-vivo* fluorescence^18,19^. Several protocols fluorescence-preserving protocols have been reported in the litterature^20,21^. We adapted the protocol developed by Zhanmu et al^20^ which replaces ethanol steps with tertiary butanol (TBA) during the dehydration process. We show that fluorescence is conserved in a mouse tumor heterogeneously expressing GFP after undergoing TBA dehydration. The sample was imaged with the SBFI approach and sections were subsequently imaged using a confocal microscope. Spectral imaging on the sections confirmed the presence of GFP-specific signal. The overlay of the images acquired by these two modalities show that GFP-specific signal is indistinguishable from the autofluorescence signal with the SBFI method (Supplemental Figure S5). Fine spectral filtering of these overlapping sources of fluorescence will be required to fully distinguish specific GFP signal from non-specific autofluorescence. This is part of the future developments of our technology. However, the possibility of recovering sections associated with the preservation of fluorescence during sample preparation can, in the current configuration of the device, allow correlative approaches between autofluorescence signal recorded with our system and the specific fluorescence signal obtained by confocal microscopy.

In this work, we show the application of the apparatus onto a microtome. However, one of the advantages of this system is its ease of integration with various types of instruments. Implementation of the system on a cryostat is highly relevant as sample freezing is more suited than paraffin embedding for certain experimental models. Indeed, preparation of frozen samples does not affect fluorescent labeling^18^ and is suitable for samples which cannot withstand dehydration and require minimal tissue processing steps. This is true for hydrogels or hydrated matrices that can not undergo dehydration without drastic morphological changes. This method could offer a new and simple alternative to image 3D bioprinted samples and matrix-embedded organoids which commonly lack sufficient transparency to be imaged in microscopy. In addition, 3D bioprinting typically relies on bioinks which are composed of constituents of the extracellular matrix, and are associated with high levels of autofluorescence (gelatin, elastin, collagen)^17^. There are different SBFI systems adapted to frozen samples and designed to recover full color images^22–24^ or fluorescence signals^25,26^. Inspired by this literature, we have tested the feasibility of using our imaging apparatus to frozen samples, by simply contrasting fluorescence tissue with OCT (Optimal cutting temperature compound) embedding, an opaque white compound^25,26^. To date, our system on the cryostat has not yet been optimized for automation, but this is part of ongoing research which will further expand the field of its applications.

This work presents a complementary SBFI methodology to the already existing approaches. Our method proves to be relevant for label free-imaging of large samples and provides a quantitative signal that can be used for further post processing analysis. Working in an open access microscopy and histology structure, the development of this system came from a real need to have an accessible and fast method allowing to combine 3D volume analysis as well as obtaining histological slices for large samples. We believe that this system could be useful in a large number of studies and can evolve according to the needs of scientific projects.

## Material and Methods

### Animals and plants

The different mouse organs collected for this study came from C57BL6/KaLwRij mice. Mice were sacrificed by cervical dislocation under isoflurane anesthesia. All mice experiments were approved by the Animal Experimentation Ethic Committee of the Pays-de-Loire (protocol no. CEEA.2013.2). Mice were housed in the UTE animal facility (University of Nantes, license no. B-44-278) under conventional conditions. Shrimp was a Neocaridina davidi adult female, recovered after a natural death and acquired from a pet store. Zebrafish was kindly provided by Jean Jacques Lareyre (INRA, UPR 1037, Rennes). Broccoli (Brassica oleracea var. italica) was purchased from a grocery store in Nantes (France). Clover (Trifolium repens) was recovered in a public garden in Nantes (France).

### Sample preparation

Samples were fixed in formol 10% for 24h to 48h depending on their size (between 0.5 and 2 cm). Samples were then dehydrated in absolute ethanol then cleared in isopropanol and finally included in ORO paraffin. To allow better paraffin penetration in the tissues, samples were let to sit for 1-2hr in warm ORO paraffin before cooling to form the blocks. For mineralized tissues like bone, a decalcification step was necessary prior to the dehydration. Samples were immersed in decalcifying solution (Decalc, Histolab, Denmark) for several days until the sample was soft enough to be cut.

### Microtome specification

For these experiments, the camera was placed in front of a microtome HM 355S (Epredia, NH, USA) with STS system and cool-cut system. Samples were cut with blades (MX35 Premier, Epredia, NH, USA). Hard or brittle samples (bones, zebrafish, liver) were cut with specific blades (#4810050, Trajan, Australia). To obtain a nearly isotropic 3D volume, the cut step was chosen according to zoom resolution (3, 5, 10 & 20 μm respectively for zoom 3, 2, 1 & no zoom).

### Camera and optical system specifications

The imaging system included a CMOS camera (Grasshopper GS3-U3-23S6M-C, FLIR). Images are acquired through a 50mm lens (Xenoplan 2.8/50, Schneider Kreuznach) and saved in 16-bit with a 1024×1024 pixels definition. Additional lenses could be used in front of the system to adapt field of view to sample size (achromatic lenses for zoom 1 AC254-150-A-ML, zoom 2 AC254-75-A-ML and zoom 3 AC254-35-A-ML, Thorlabs). Excitation light is provided by a fibered 470 nm LED source (Mightex). A dichroic beam splitter (DMLP490R, Thorlabs) and an emission filter (FF01-496, Semrock) are used to recover emitted photons with a wavelength greater than 500 nm. LED power was measured with an optical power meter PM100D (Thorlabs, Maison Lafitte, France).

The system is able to trigger a microtome cutting cycle using its footswitch input and mimicking pedal activation by the user. Two images are sequentially recorded at each acquisition. The first one is the fluorescence image with the excitation light on. The second image produced has the same parameters except that the excitation light is turned off. It effectively records the acquisition background noise. A subtracted image of fluorescence signal minus noise is generated by the software in real-time.

### 3D volume generation

The series of subtracted images were saved as a stack with Fiji software^27^ and aligned with the Stackreg plugin^28^ (translation registration only) when it was necessary. 3D visualization was done with the plugin ClearVolume in Fiji for the figure 2h. Figure 6c and 7c was done with Arivis software.

### Image analysis

The 3D segmentation of the mouse liver and its vascularization presented in Figure 6(a-c) was done on Fiji software. Liver volume and blood vessels were segmented using default thresholds and volumes were determined with the 3D Objects Counter. The vascular network was reconstructed using the Skeletonize plugin and characterized with Analyze skeleton.

Data presented in Figure 7(a,b,d) were correlated using eC-CLEM^29^ in Icy^30^. The slice number in the SBFI system was recorded when sections were kept, and further stained for HPS. The registration was then performed rigidly in eC-clem between the 2D slice from the 3D SBFI corresponding to the stained section number, by identifying 14 common landmarks points. Regions of interest were drawn manually in Icy as 2D Polygon on the HPS stained section, and then copy and pasted directly on the registered 2D SBFI section. The reverse transformation was applied to the 2D stained section and placed in the original SFBI volume at its correct slice position in order to visualize the HPS stained section on the 3D volume. Heart was segmented on the SFBI stack using ITK-SNAP^31^ by the semi-automatic segmentation snake based method, followed by a manual correction for the ventricles segmentation. The generated 3D binary mask was then imported in Icy, and a 3D ROI was generated, allowing direct quantification of the 3D ROI volume. Visualization was performed under Icy with volume slicing in the 3D viewer.

### Histological staining

After dewaxing and rehydration of sections, HPS (Hematoxylin (Diapath, Italy) Phloxin (Biognost, Croatia) and Saffron (VWR)) staining was conducted according to manufacturer protocols. This staining allows visualization of nucleus in violet, cytoplasm in pink and collagen in orange. Slides were then scanned using a slide scanner (Nanozoomer, Hamamatsu Photonics, Japan) with x20 magnification.

## Data availability

The datasets generated and/or analyzed during the current study are available from the corresponding author on reasonable request.

## Acknowledgements

We thank Justine Perrin (UMR U1307-CRCI2NA, Nantes), Malo Daniel (UMR U1064-CRTI, Nantes), Tacien Petithomme (UMR U1307-CRCI2NA, Nantes), Gwennan André-Grégoire (UMR U1307-CRCI2NA, Nantes), Xavier Druart (PRC, UMR INRAE CNRS; PIXANIM INRAE, Nouzilly) and Jean Jacques Lareyre (INRA, UPR 1037, Rennes) who kindly donated the samples presented in this study. We also thank Jean Mérot, Constance Delwarde and Pascal Aumond (UMR 1087, Institut du Thorax, Nantes) for the fruitful discussions on this system development. We acknowledge the IBISA MicroPICell facility (Biogenouest), member of the national infrastructure France-Bioimaging supported by the French national research agency (ANR-10-INBS-04). Particularly, we thank Annabelle Justin for her assistance on sample preparation and HPS staining and Steven Nedellec for his revision of the manuscript. We acknowledge ANR-18-CE45-0015 and Biogenouest for financial support.

## Author Contributions

M.M. and P.d.V. prepared the samples, designed and conducted the experiments and drafted the manuscript. P.H. and P.D. designed and constructed the imaging system, P.H characterized the system. M.F. analyzed liver vascularization. S.B. optimized the preservation of fluorescence protocol during sample preparation. G.S. developed the acquisition software for automatization. P.P.G. carried out the correlative analysis on the mouse embryo experiment, analyzed embryonic heart segmentation and supervised the project. All of the authors revised the manuscript critically for important intellectual content, gave final approval of the version to be published and agreed to be accountable for all aspects of the work.

