## Supplementary methods and figures for "Label-free imaging of large samples: 3D rendering and morphological analysis within histological workflows using serial block face imaging"

### Supplementary Information

#### Supplementary Methods

*SBFI system adapted to frozen sample*

Right after organ harvesting, samples were embedded in OCT (Tissue freezing medium, MM France) and frozen by immersion in isopentane cooled with liquid nitrogen. Frozen samples were kept at -80°C until they were processed. The camera was positioned in front of a cryostat (CM1950, Leica Biosystems, Nussloch, Germany).

*Fluorescence signal preservation during paraffin embedding*

Subcutaneous mouse tumor injected with adeno-associated virus (AAV) expressing GFP was fixed with formol 10% for 24h and dehydrated with optimized Tert-butanol (TBA) protocol inspired from Zhanmu et al[21]. Sample was successively immersed in solutions of TBA 75%, TBA 95% and two baths of TBA 100%, for 24 hours each. Then the sample was embedded in ORO paraffin and image with SBFI. Individual sections were collected and imaged on a confocal microscope (A1 RSi, Nikon Instruments Inc., Japan) at 488 nm.

#### Supplementary Figures

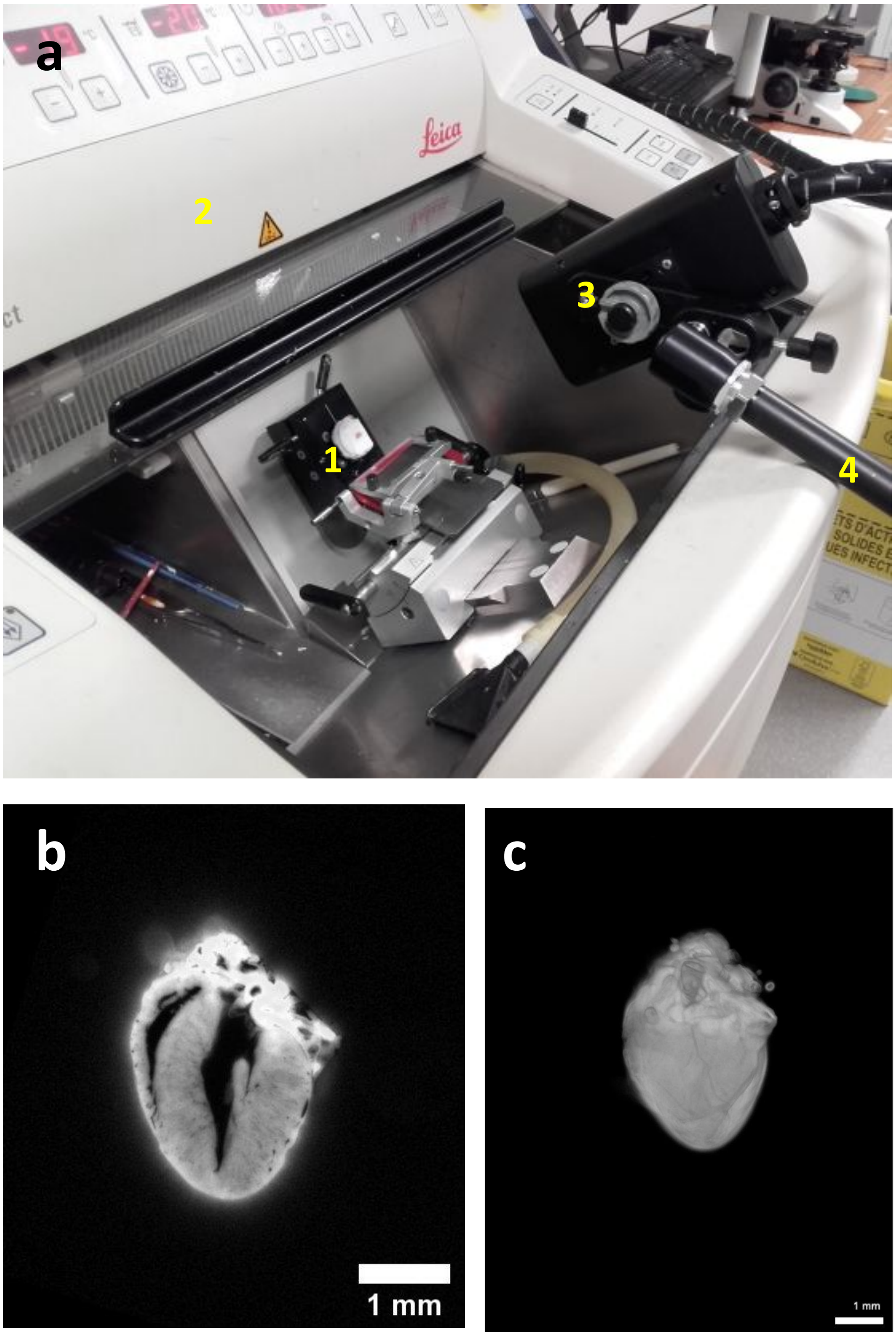


**Supplemental Figure S1:** The imaging system was mounted on a cryostat (CM 1950, Leica). **(a)** Image of the system assembly. A frozen samples embedded in OCT medium (1) is placed in the cooled chamber of the cryostat (2). After each cut, an image of the surface of the block is acquired by the camera (3) placed orthogonally to the surface of the block using an adjustable arm (4). The camera was connected to the control unit which itself is connected to a laptop computer (not shown). Image capture is done manually via interface. A typical autofluorescence image of a frozen mouse heart embedded in OCT **(b)** and the 3D visualisation of the sample with Arivis software **(c).**

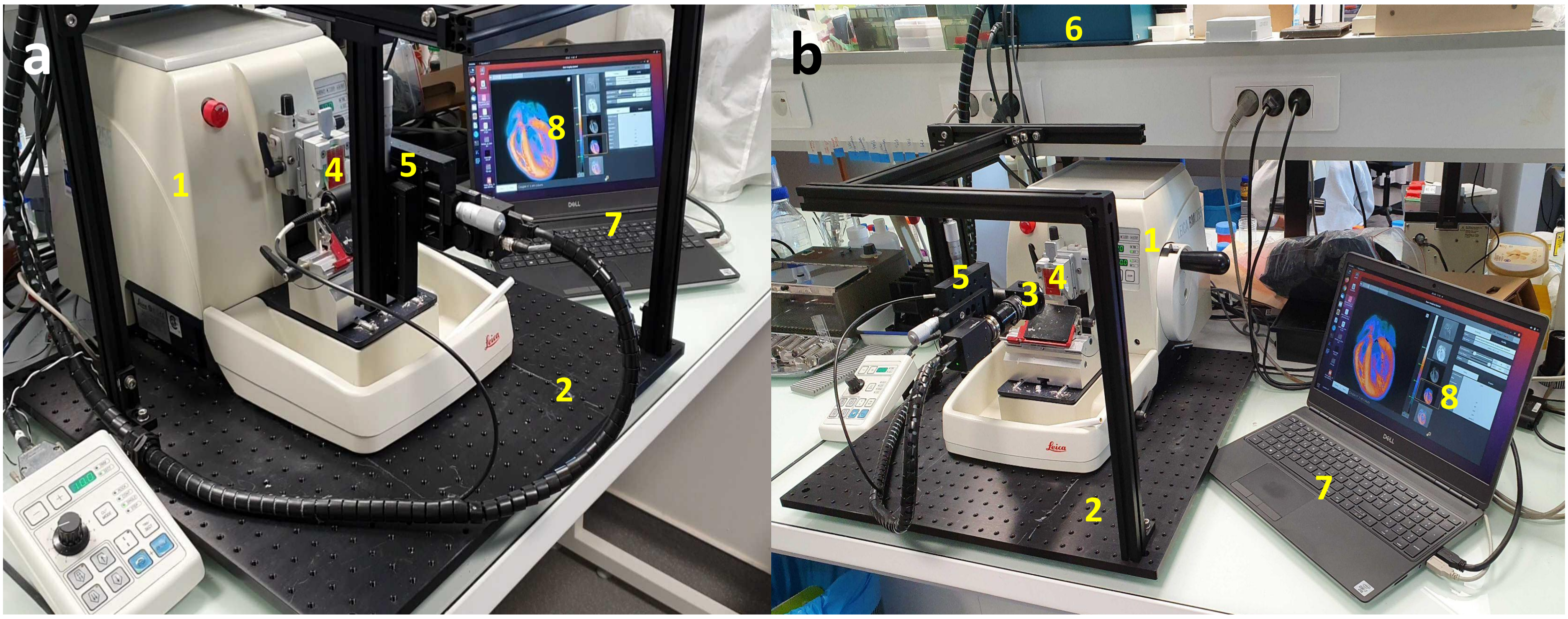


**Supplemental Figure S2 :** SBFI using an automatic rotary microtome (RM2255, Leica). Whole view images of the initial imaging system **(a-b)**. The rotary microtome (1) is placed on an optical table to which is attached an aluminum frame (2) that holds the camera (3) orthogonally to the surface of the mounted sample (4). The camera is attached to a Y and Z positioning stages (5). The microtome is connected to the control unit (6) which itself is connected to a laptop computer (7). The acquisitions are controlled by the control software (8) that triggers the microtome through the control unit.

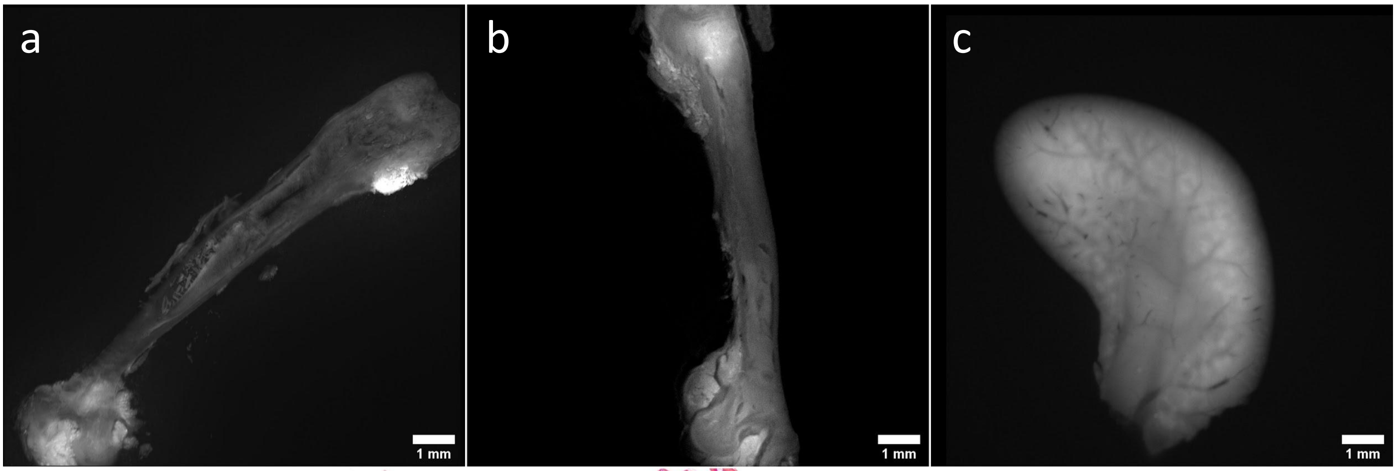


**Supplemental Figure S3:** Autofluorescence images obtained during SBFI acquisitions, corresponding to recovered sections of mouse tibia (a,b) and a mouse liver (c) shown in Figure 5.

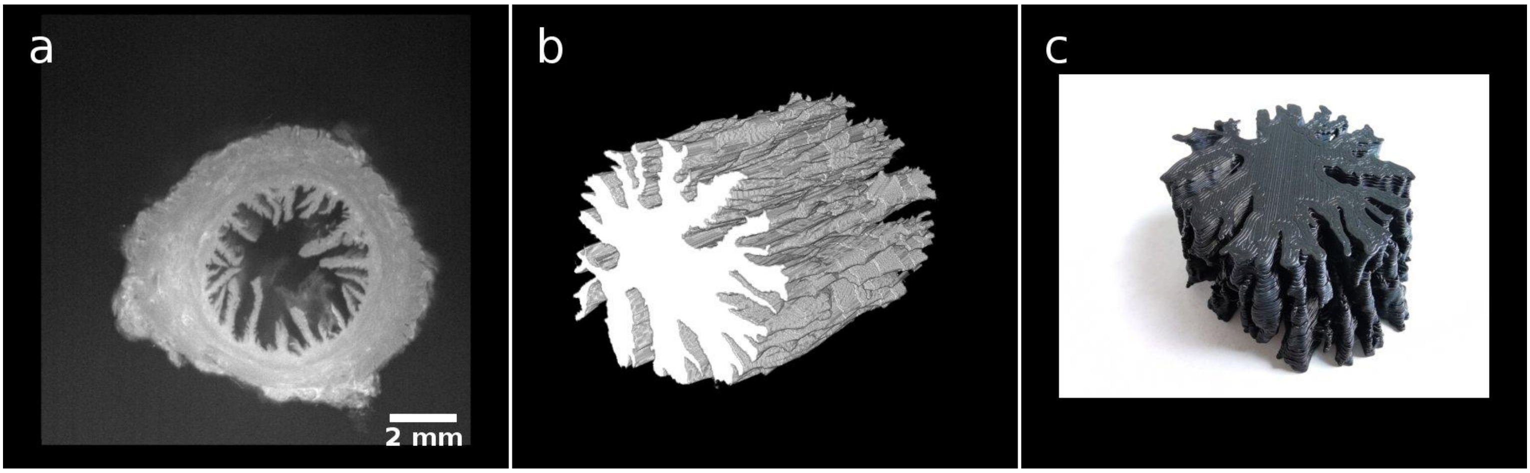


**Supplemental Figure S4:** 3D printing from a segmented structure. A segment of a ewe cervix was processed (a). Segmentation of the lumen on the image stack was performed to reveal cervical internal structures and organisation using Fiji (b). A mesh was generated from the segmented object using Opencv and was 3D printed using a  Ender 3 V2 3D printer (Creality, China)(c).

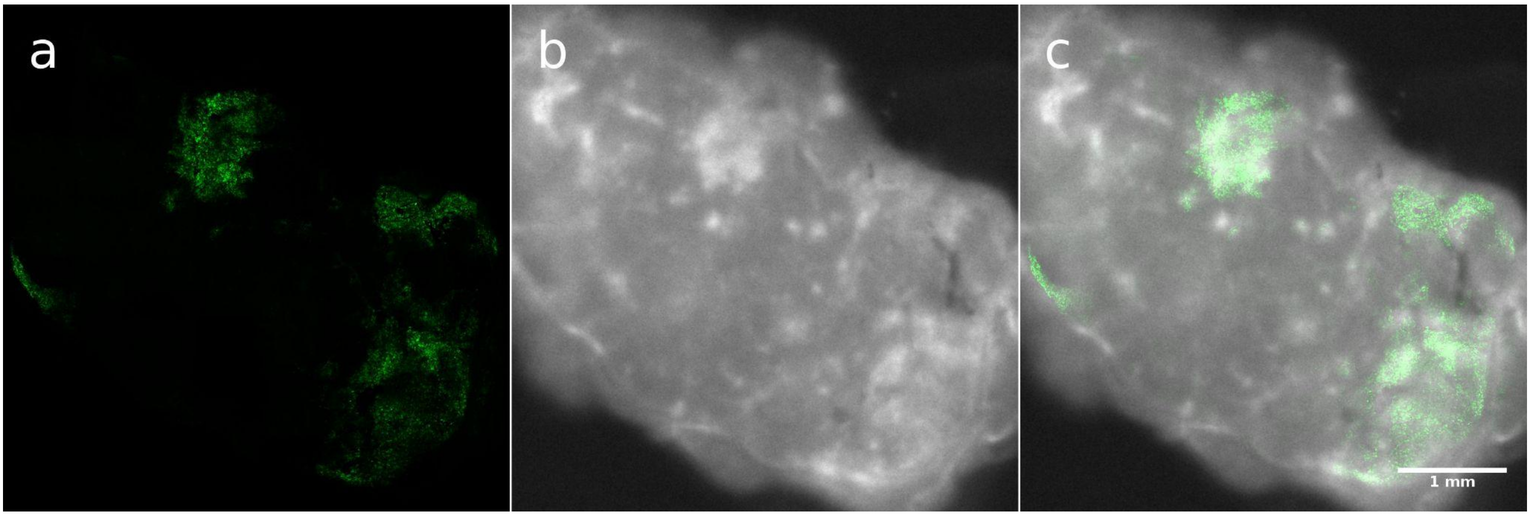


**Supplemental Figure S5:** Validation of a fluorescence preserving dehydration protocol on a heterogeneously GFP-stained mouse tumor. **(a)** GFP signal of the mouse tumor detected at 488nm using a confocal microscope. **(b)** Autofluorescence image of the tumor acquired with the SBFI system. The overlay of the two imaging modalities is shown in **(c)**.
